# Repeated mild traumatic brain injury does not affect sleep or epileptiform activity one-month post-injury in a knock-in mouse model of Alzheimer’s disease

**DOI:** 10.64898/2026.07.24.740449

**Authors:** Victoria Carriquiriborde, Jefferey Yue, Wai Hang Cheng, Taha Yildirim, Jianjia Fan, Sean Tok, Michael Kelly, Cheryl L. Wellington, Brianne A. Kent

**Affiliations:** Institute for Neuroscience and Neurotechnology, Simon Fraser University, 8888 University Drive, Burnaby, BC, Canada, V5A 1S6; Department of Molecular Biology and Biochemistry, Simon Fraser University, 8888 University Drive, Burnaby, BC, Canada, V5A 1S6; Department of Psychology, Simon Fraser University, 8888 University Drive, Burnaby, BC, Canada, V5A 1S6; Department of Pathology and Laboratory Medicine, University of British Columbia, 2215 Wesbrook Mall, Vancouver, BC, Canada, V6T 1Z3; Djavad Mowafaghian Centre for Brain Health, University of British Columbia, 2215 Wesbrook Mall, Vancouver, BC V6T 1Z3; School of Biomedical Engineering, University of British Columbia, 2215 Wesbrook Mall, Vancouver, BC V6T 1Z3; International Collaboration on Repair Discoveries, Blusson Spinal Cord Centre, Vancouver General Hospital, 818 W 10th Ave, Vancouver, BC V5Z 1N1

**Keywords:** Concussion, Alzheimer’s disease, EEG, sleep, seizure, amyloid

## Abstract

Traumatic brain injuries (TBIs) are associated with increased risk of neurodegenerative disease, including Alzheimer’s disease (AD); however, the mechanisms by which TBI promotes AD pathogenesis remain poorly understood. It also remains unclear whether post-TBI sequelae, including sleep disturbances and seizures, play a role in driving disease progression. To investigate these relationships, we employed a translational approach using the Closed-Head Injury Model of Engineered Rotational Acceleration (CHIMERA) of repeated mild TBI (rmTBI) and an AD knock-in mouse model to assess sleep, power spectral density, epileptiform activity, and Aβ pathology one month post-injury. RmTBI caused elevated neurofilament-light and glial fibrillary acidic protein, markers of neuronal damage. Sex differences were observed in acute injury outcomes, sleep measures, and Aβ plaque size. Specifically, females exhibited longer recovery post-injury, higher mortality, decreased non-rapid eye movement sleep duration, and larger average plaque size than males at equivalent impact energy. These findings highlight the importance of including both sexes when establishing injury severity thresholds. Future studies should incorporate validated TBI biomarkers of neural injury to define equivalent injury parameters across sexes and examine the chronic effects of rmTBI on sleep, epileptiform activity and AD pathology.

## Introduction

Traumatic brain injuries (TBI) are a major cause of disability and death worldwide (James et al., 2019), and an important public health issue. Approximately 70-80% of TBI cases are classified as mild (Leo & McCrea, 2016) and symptoms resolve within days to weeks (Willer & Leddy, 2006); however, repeated injuries can synergize and potentially lead to more profound consequences (Guskiewicz et al., 2003; Montenigro et al., 2017). Repeated mild TBI (rmTBI) are associated with increased risk of neurodegenerative disease, such as Chronic Traumatic Encephalopathy (Cherry et al., 2020; Jun et al., 2026) and potentially Alzheimer’s disease (Alanazi et al., 2024; Batty et al., 2023; Graham et al., 2022; Graham et al., 2025). RmTBI cases are most prevalent in contact sport athletes, military personnel, survivors of intimate partner violence, and people experiencing homelessness (Esopenko et al., 2024; CDC, n.d.; Janković & Pilipović, 2023; O’Connor et al., 2022; Stubbs et al., 2020).

TBI can increase pathological hallmarks of AD, including Aβ deposition, tau phosphorylation, and microgliosis (Sivanandam & Thakur, 2012). Clinical studies have found an increase in Aβ deposition following TBI, measured with positron emission tomography (PET), although the spatial distribution differs from that observed in typical AD (Hong et al., 2014; Scott et al., 2016). Post-mortem studies of head injury have also shown evidence of amyloid precursor protein (APP) accumulation and co-localization of APP with Aβ (Uryu et al., 2007). In addition, rmTBI have been associated with increased Aβ accumulation in transgenic animal models of AD (Cheng et al., 2018; Conte et al., 2004; Uryu et al., 2002). Together, these shared pathological features may suggest overlapping mechanisms linking rmTBI and AD.

RmTBI have also been shown to affect sleep quantity and quality (Alosco et al., 2021; Bryan, 2013; Sandsmark et al., 2017), with individuals reporting difficulty initiating sleep, nocturnal awakenings, and excessive daytime sleepiness or frequent napping (Sandsmark et al., 2017; Wickwire et al., 2016). These symptoms are similar to those reported in individuals with AD (Brzecka et al., 2018; Most et al., 2012). Quantitative electroencephalographic (qEEG) power spectral density (PSD) analysis provides a measure of brain oscillatory activity at each frequency, with distinct spectral profiles characterizing the vigilance states: non-rapid eye movement (NREM) sleep is dominated by delta-band power (0.1–4 Hz), rapid eye movement (REM) sleep by theta-band activity (4–8 Hz), and wakefulness by a broadband distribution with relatively greater high-frequency power. Post-mTBI alterations in PSD have been documented in both humans and rodent models (Coyle et al., 2025; Lewine et al., 2019; Napoli et al., 2012; Paterno et al., 2016). This is of particular translational relevance given the evidence of PSD shifts in AD (Kent et al., 2021, 2022) and association with glymphatic function (Dagum et al., 2025; Hablitz et al., 2019a), a system responsible for clearance of metabolic waste products including Aβ and tau (Iliff et al., 2014; Jessen et al., 2015). As such, PSD alterations may serve as markers of impaired Aβ clearance and disrupted sleep, potentially contributing to AD-related pathology following repeated head injury.

Additionally, TBI is associated with an increased risk of post-traumatic epilepsy (PTE), which may manifest as overt post-traumatic seizures or subclinical epileptiform activity (Kim et al., 2018; Lucke-Wold et al., 2015; Ronne-Engstrom & Winkler, 2006). Although PTE occurs at a lower incidence following mild TBI, the risk is compounded by repeated injury (Lolk et al., 2021; Lucke-Wold et al., 2015). Preclinical evidence from wild-type mouse models of rmTBI corroborates human findings, implicating reactive astrogliosis and oxidative stress as putative mechanistic drivers (Bugay et al., 2020; MacMullin et al., 2020; Shandra et al., 2019). Critically, PTE may not only co-occur with AD pathology but actively accelerate it, underscoring the clinical relevance of characterizing epileptiform activity in the context of rmTBI.

Building on our previous work in the APP/PS1 transgenic mouse model (Yue et al., 2026), here we employ the APP^NL-F^ knock-in model, which avoids the confound of APP overexpression (Saito et al., 2014; Sasaguri et al., 2017) and better recapitulates human amyloid pathology (Saito et al., 2014). Injuries were delivered at six months of age, coinciding with the onset of Aβ accumulation in this mouse model, using the Closed-Head Injury Model of Engineered Rotational Acceleration (CHIMERA), a platform validated for its ability to replicate the diffuse biomechanics of human mTBI (Namjoshi et al., 2014). Consistent with our prior APP/PS1 work (Yue et al., 2026), outcomes were assessed one month post-injury, a subacute timepoint that remains understudied. Sleep architecture, PSD, and epileptiform activity (EA) were evaluated via EEG/EMG recordings, and protein markers of neuronal damage and amyloid pathology were quantified by immunoassay. We hypothesized that rmTBI would cause sleep impairments, reductions in slow-wave sleep, and increases in EA, plaque burden, and neuronal damage at this timepoint. Contrary to expectations, no changes in sleep, PSD, EA, or plaque burden were observed one month post-rmTBI, despite elevated neurofilament-light (NF-L) and glial fibrillary acidic protein (GFAP). Notably, female mice exhibited greater susceptibility to injury-related adverse outcomes following the third rmTBI.

## Methods

### Animals

All procedures in this study were approved by the Simon Fraser University Animal Ethics Committee (Protocol 1330P-21) and followed the guidelines from the Canadian Council on Animal Care. This study adheres to the Animal Research: Reporting of In Vivo Experiments (ARRIVE) guidelines for reporting animal research. Figure 1 illustrates the study protocol. Six-month-old homozygous APP^NL-F^ mice (Saito et al., 2014) containing the humanized Aβ sequence, the Swedish mutation (KM670/671NL) and the Beyreuther/Iberian mutation (I716F) on a C57Bl/6J background [C57BL/6-App<tm2(NL-F)Tcs>/TcsRbrc (RBRC06343)], were used in this study. Mice were bred in-house in the Wellington laboratory (University of British Columbia). We chose this amyloid precursor protein (APP) knock-in (KI) model of AD because it avoids artifacts related to the overexpression of APP present in other mouse models of AD. Aβ plaque accumulation in this mouse model starts at 6 months of age and increases exponentially starting at 12 months of age. All animals were housed in plastic clear cages under 12:12 light:dark cycle. Both sexes were grouped housed before the study procedures started and singled housed thereafter. Food and water were available *ad libitum*.

**Figure 1.**
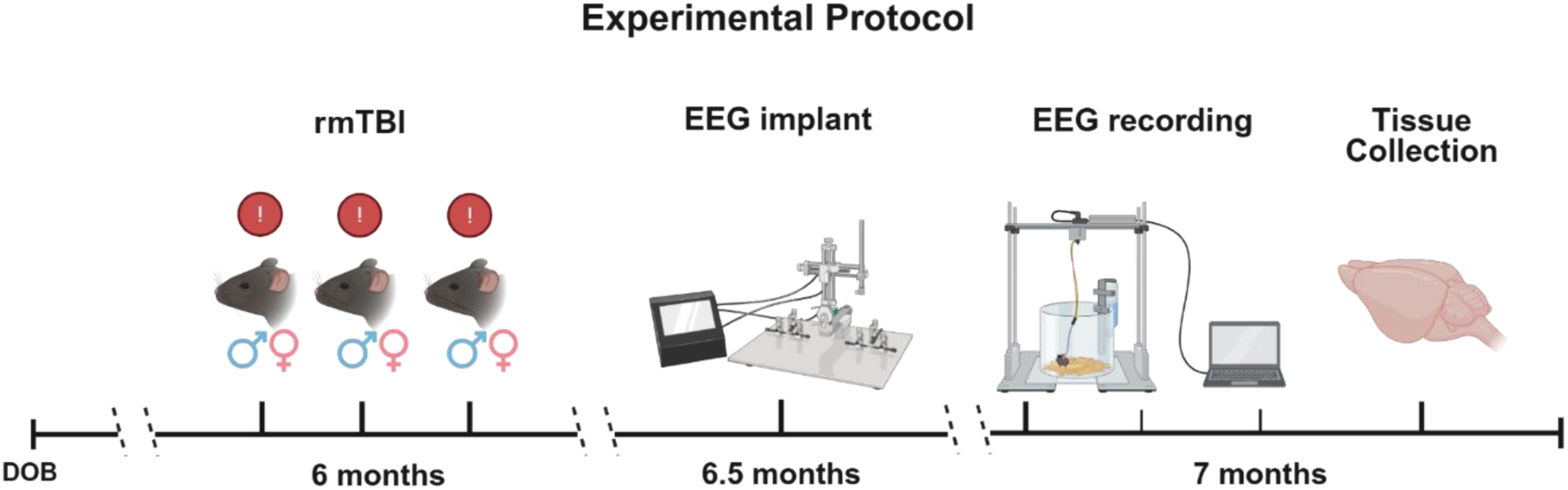
Schematic diagram illustrating the experimental protocol. At six months of age APPNL-F mice received 3 consecutive mTBIs 24 h apart. After recovering, mice received EEG/EMG intracranial implantation. At 7 months of age, the mice underwent 72 h of EEG/EMG recording. Tissue collection was completed immediately after recording. Created in BioRender https://BioRender.com/6h4as63

#### 1. Repeated mild TBI (CHIMERA)

CHIMERA procedures followed the protocol previously described (Yue et al., 2026). Briefly, mice received meloxicam subcutaneously (SQ, 1mg/kg) 30 min before being restrained on the CHIMERA device. Mice were anesthetized with isoflurane (0-5%), received saline (SQ, 0.5mL) for hydration, and eye gel was applied. Total duration of isoflurane exposure was 4 - 8min. Once the animal was positioned as specified in Namjoshi et al. (2014), the impact was delivered covering a 5 mm diameter surrounding bregma. Mice received the CHIMERA impact at 6 months of age. Both males and females received 3 TBI at 0.7J, each TBI was 24 h apart. Sham animals underwent the same procedures except for the piston impact. Mice recovered in a warmed cage supplemented with hydrogel.

#### 2. Loss or Righting Reflex and Weight

The loss of righting reflex (LRR) was calculated as the time interval from the termination of isoflurane to the successful attempt of righting for each mouse after injury. In TBI research, LRR is used as an analog of loss of consciousness in humans (Berman et al., 2023). Longer LRR times are associated with cognitive and memory impairments and worse neurological outcomes (Berman et al., 2023). Body weights were measured immediately before the first TBI (baseline) and 6 hours after each impact (post-mTBI).

#### 3. Electrode implantation

2EEG/1EMG headmount (catalog number 8201-SS; Pinnacle Technology, Lawrence, KS, USA) implant surgery was performed two weeks after the CHIMERA protocol. Briefly, four stainless steel screw electrodes were inserted into the skull at coordinates relative to bregma: AP +/- 3mm, ML +/- 1.5mm. EMG wires were inserted into the nuchal muscle. Post-operatively, lactated ringer (SQ, 5mg/kg) was provided for hydration, and buprenorphine (0.05mg/kg) was provided for 3 days for pain relief. Mice were allowed to recover from surgery for two weeks before EEG recordings started. Surgical procedures have been described in detail previously (Yue et al., 2026).

#### 4. Electroencephalography recording

EEG/EMG recordings were collected using a tethered system (8200-K1, Pinnacle Technology) and Sirenia® Acquisition software (v2.2.1). Sample rate was 400Hz, preamp gain was 100dB, and low-pass filter was 40Hz for EEG and 100Hz for EMG. Each mouse was recorded for 72 consecutive hours approximately one-month post-TBI. Animals were euthanized after the 72 h recording, brain tissue was collected and processed.

#### 5. Assessment of vigilance state

The assessment of vigilance state was performed using the same parameters and scoring criteria as described previously by Yue et al. (2026). The three vigilance states (NREM, REM and wake) were scored using the Sirenia Sleep Pro software (v3.0.1). Only the last 24 h of the recordings were scored and used for analyses, to allow for habituation of animals to the new cage and tethered system in the first 48 h.

#### 6. Power Spectral Density Analysis

Power spectral density (PSD) provides a quantitative measure of how neural activity is distributed across EEG frequencies. We calculated power at each frequency and each electrode separately as described previously by Yue et al. (2026). Epochs containing artifacts were removed from the data using a thresholding method; specifically, epochs were excluded if the power was above the 90^th^ percentile for that animal within that vigilance state. Once data were artifact-free, relative power was calculated by dividing the absolute power at each frequency by the total power across frequencies (0.5-30Hz). We analyzed the delta (0.5-4Hz), theta (4-8Hz), alpha (8-13Hz) and beta (13-30Hz) bands (Hablitz et al., 2019b; Kadam et al., 2017; Kent et al., 2018).

#### 7. Epileptiform Activity Analysis

Epileptiform activity (EA) detection was performed using a custom Matlab (R2023a, v9.14) script, available on GitHub (https://github.com/Translational-Neuroscience-Lab/TBI_project). Continuous EEG recordings were segmented into 10-s epochs and assigned a vigilance state (NREM, REM, or wake) based on the scoring. For each epoch, the signal envelope was computed using the Hilbert transform, and a detection threshold was set at 6 standard deviations (SD) above the envelope mean, in accordance with recommendations for preclinical epileptiform spike detection (Jin et al., 2022). Epochs exceeding this threshold were flagged and subsequently reviewed using an integrated visualization tool, through which a researcher blinded to the treatment condition classified each event as a single spike, poly spike, or seizure. Only the final 24 h of each recording were included in EA analyses.

A single spike was defined as a cortical epileptiform discharge with an amplitude exceeding 6 SD of the epoch envelope mean; events attributable to movement artifacts were excluded during the visual review. Poly-spikes were defined as two or more consecutive spikes meeting the same amplitude criterion. Seizures were defined as high-amplitude, high-frequency discharges lasting more than 10 s. To account for differences in time spent in each vigilance state across animals, EA counts were normalized to the total hours spent in each respective state.

#### 8. Biosample collection

Mice were deeply anesthetized by isoflurane (5% flow rate) then euthanized with CO_2_ and blood samples collected via cardiac puncture and mixed with EDTA to prevent coagulation. Plasma was extracted after centrifugation and stored at -80°C until analysis. Mice were perfused and brain tissue was extracted. The brain was bisected across the midline, with one hemibrain snap-frozen on dry ice and kept in -80°C freezer, and the other hemibrain fixed in 4% paraformaldehyde for 3 days and then cryoprotected in 30% sucrose. The cryopreserved hemibrain was coronally sectioned at 40μm using cryotome (Leica).

Hemibrain lysate was generated in serial extraction as previously described (Robert et al., 2016). Frozen hemibrain was ground, then homogenized in four volumes of carbonic buffer (100 mM Na2CO3 and 50 mM NaCl, pH 11.5) supplemented with 1mM phenylmethanesulfonyl fluoride (PMSF) and protease inhibitor cocktail (Roche, cat #11836170001), then sonicated in 20% output for 10s. After centrifugation, the supernatant was collected and neutralized with 1M Tris-HCl (pH 6.8) to yield a final pH of ∼7.4. The pellet was resuspended in four volumes of 5M guanidine hydrochloride (5 M GuHCl, 50 mM Tris-HCl) and incubated at room temperature with rotation for 3h. Both fractions were stored at -80°C until analysis.

#### 9. Plasma Biomarker Analysis

Plasma levels of murine glial fibrillary acidic protein (GFAP) were quantified by a customized immunoassay (Yue et al., 2026) using a pair of rabbit antibodies against mouse GFAP. Plasma was diluted 4 times with PBS containing 1% bovine serum albumin (Sigma Aldrich, cat. # A7906). Plasma levels of murine neurofilament-light (NF-L) were measured by the MesoScale Discovery (MSD) Neurofilament-light assay kit (Mesoscale, cat. # K1517XR-2) following the manufacturer’s instructions. Plasma was diluted 6 times with assay diluent provided by the kit. The researcher was blinded to the experimental conditions. All samples with reported comparisons were analyzed within the same run.

#### 10. Immunohistochemistry

Immunohistochemistry of amyloid plaques was performed as previously described (Yue et al., 2026) to examine cortical amyloid plaques in the temporal isocortex (ISOCTX) defined according to anatomical landmarks illustrated on the Allen mouse brain atlas (https://mouse.brain-map.org/static/atlas). Every tenth coronal brain section at 40 µm-thickness was sampled, and up to three sections per region were analyzed. After digesting sections in 88% formic acid for 5 mins and blocking in 5% milk diluted in 0.25% Triton X-100, sections were incubated with anti-Aβ (6E10; BioLegend, cat # 803002) antibody overnight, then incubated with biotinylated secondary antibody (SouthernBiotech, cat # 1072-08). Sections were further developed using the ABC system (Vector Lab, cat # PK-6100) then chromogen development using 3,3’-Diaminobenzidine (DAB). Mounted sections were dehydrated and coverslipped, then imaged using an Axio Scan.Z1 slide scanner (Zeiss) at 20x magnification, followed by resizing at 30% and image optimization using the Zen software (v3.7).

#### 11. Aβ_42_ ELISA

Soluble (carbonic fraction) Aβ_42_ in hemibrain lysates were quantified using a commercial ELISA kit (WAKO, cat # 296-64401) according to the manufacturer’s instructions. The carbonic fraction was diluted 1:10 with assay buffer. Every sample was loaded in triplicate, and the average value of the three readings was reported.

#### 12. Statistical Analysis

Data normality was assessed using the Shapiro-Wilk test (Mishra et al., 2019). LRR and body weight data were analyzed using a mixed-effects model with the Geisser-Greenhouse correction, comparing four groups (male sham, female sham, male TBI, and female TBI) across three timepoints (1^st^, 2^nd^, and 3^rd^ TBI) for LRR, and four timepoints (baseline and 3 TBI) for body weight, with Tukey’s correction applied for post-hoc multiple comparisons. Given a violation of the normality assumption, the relationship between LRR and body weight within the rmTBI group was examined using Spearman rank correlation, computed separately at each timepoint. Vigilance state durations, state transition counts, and light/dark period were analyzed using mixed-effects model with Geisser-Greenhouse correction. The vigilance state data were analyzed using a two-way ANOVA to compare sex and grouping effects. GraphPad (v10.3.1) was used for all vigilance state, LRR, and body weight analyses.

Normalized EEG power spectral density (PSD) data were analyzed using mean-centered partial least squares (PLS) in MATLAB (R2023a, v9.14) (McIntosh & Lobaugh, 2004). PLS is a multivariate method well-suited for neuroimaging data in which dependent measures are highly collinear, as it identifies latent variables (LVs) capturing the relationship between experimental conditions and patterns of spectral power across the full frequency spectrum in a single analytical step, circumventing the need for multiple comparisons (Cragg et al., 2011). Analysis structure and procedure followed those described in Cragg et al. (2011). Separate LVs were derived for each channel (frontal and parietal) and vigilance state, with statistical significance assessed via permutation testing and bootstrap resampling.

EA data were compared between treatment groups (sham vs. TBI) using Mann-Whitney U tests. The interaction between vigilance state and treatment group on EA was examined using mixed-effects model with the Geisser-Greenhouse correction. A secondary exploratory analysis to assess sex effects was conducted using two-way ANOVA for single spikes and poly-spikes, and a mixed-effects model with the Geisser-Greenhouse correction for epileptiform activity over vigilance states separating treatment groups by sex.

Plasma NF-L and GFAP levels were compared between treatment groups using Mann-Whitney U tests, with sex assumed not to influence baseline biomarker concentrations. Amyloid plaque burden, quantified as the percentage of immunoreactive area within the isocortex (ISOCTX) using ImageJ, and Aβ_42_ ELISA data were analyzed using two-way ANOVA with treatment group and sex as independent factors, performed in GraphPad Prism.

All statistical tests were two-tailed with significance set at p ≤ 0.05.

## Results

### Female mice experienced higher morbidity and mortality following rmTBI

The initial sample comprised n=20 animals, with equal number of males and females. All animals received three impacts at 24-h intervals at an impact energy of 0.7J. No complications were observed in males rmTBI animals, all of whom recovered fully following each impact. In contrast, two female rmTBI animals required euthanasia due to poor post-injury recovery, one prior to and one following the third impact. These animals were included in the loss of righting reflex (LRR) and body weight analyses but excluded from all subsequent assessments.

Animals in the rmTBI group exhibited a significantly prolonged LRR compared to sham controls. Sham animals of both sexes recovered the prone position within an average of 1.3 min following anesthesia, whereas male and female rmTBI animals required an average of 4.2 and 9.1 minutes, respectively. Mixed-effects model analysis comparing LRR across the four groups (male sham, female sham, male rmTBI, and female rmTBI) and three timepoints revealed a significant group x time interaction (p<0.05; **Fig S1A**). Post-hoc comparison using Tukey’s correction identified significantly longer LRR in male rmTBI relative to male sham animals across all three timepoints (p<0.05), and in female rmTBI relative to female sham animals following the 1^st^ and 3^rd^ impacts (p<0.05), but not the 2^nd^.

We hypothesized that the smaller body mass of female mice contributed to the elevated morbidity and mortality observed in the female rmTBI group. Mixed-effects model analysis revealed significant main effects of group (p<0.001) and time (p<0.001) on body weight, with no significant group x time interaction (p>0.05; **Fig. S1B**). Female rmTBI animals had lower weight than both male groups (sham and rmTBI) at all timepoints (p<0.05), but did not differ from female sham animals. The two female rmTBI animals that were euthanized prior to EEG/EMG recordings were retained in the body weight analysis, as their lower body mass may have contributed to their poor recovery following the second and third impacts.

Spearman rank correlation analysis revealed a significant negative relationship between body weight and LRR in the rmTBI group at the second (r=-0.64, p<0.05) and third (r=-0.79, p<0.05) impacts, indicating that lighter animals exhibited prolonged recovery times. No significant correlation was observed at the first impact (**Fig. S1 C-E**).

### Sleep was not impacted by rmTBI at one month post-TBI

Mixed-effects model revealed no significant differences in the total time spent in wake, NREM, or REM sleep between sham and rmTBI groups and no significant interaction group x vigilance state (p>0.05; **Fig. 2A-C**). Analysis of lights-on and lights-off periods confirmed the expected main effect of light period for all three vigilance states, with no significant main effect of treatment group or group x vigilance state interaction (p>0.05; **Fig 2E-G**). State transition counts, a measure of sleep fragmentation, did not differ significantly between groups (p>0.05; **Fig 2D**), nor was a significant interaction effect detected between groups and vigilance state transitions (p>0.05).

**Figure 2.**
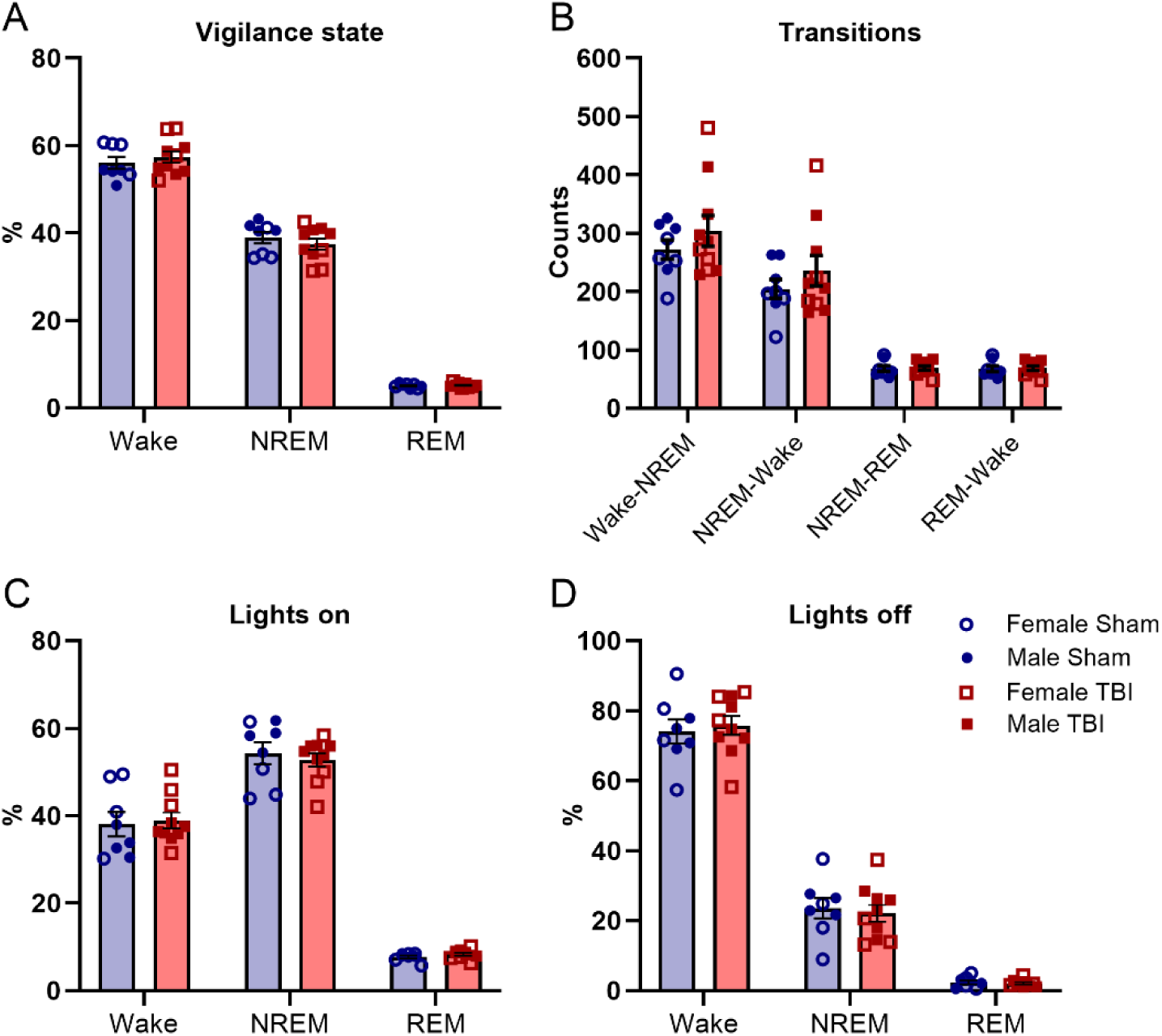
Sleep architecture of APPNL-F mice, one month after rmTBI. **(A)** Percentage of time spent in NREM, REM, and wake. **(B)** Transition counts between vigilance states wake-NREM, NREM-wake, NREM-REM, REM-wake. **(C)** Percentage of time spent in each vigilance state during lights on and **(D)** lights off. All data are expressed in mean ±SEM and analyzed by mixed-effects model with the Geisser-Greenhouse correction. Sham data are shown in blue, and rmTBI data are down in red. Females are shown with open symbols and males are shown with solid symbols. Each dot represents the average for a single mouse. Male TBI n=6, male sham n=4; female TBI n=4, female sham n=4.

Two-way ANOVA with sex and treatment group as independent factors revealed significant main effects of sex on wake (p<0.05) and NREM (p<0.05) duration during the lights-on period, with no significant sex differences observed during REM sleep or the lights-off period. Female animals spent more time in wake and less time in NREM relative to males during lights-on period, independent of treatment group (**Fig. S2**).

### No shift in power spectral density one month post-TBI

Mean-centered partial least squares (PLS) analysis revealed no significant differences in PSD distribution between sham and rmTBI groups at either the parietal (**Fig. 3**) or frontal (**Fig. S3**) electrodes across any vigilance state (wake, NREM, REM) or light periods. Due to poor EEG signal quality, one female sham mouse was excluded from the parietal PSD analysis, and one male TBI mouse was excluded from the frontal PSD analysis. A secondary exploratory analysis examining sex as a factor identified significant differences in frontal PSD during wake, at both lights-on (p < 0.005) and lights-off (p < 0.05) periods (**Fig. S4 A-B**). Female mice exhibited greater power in the slow oscillation range (< 1 Hz) and reduced power in the theta rage (4-6 Hz) relative to males, independent of treatment group, only during wake. There were no sex differences in the other states (NREM and REM).

**Figure 3.**
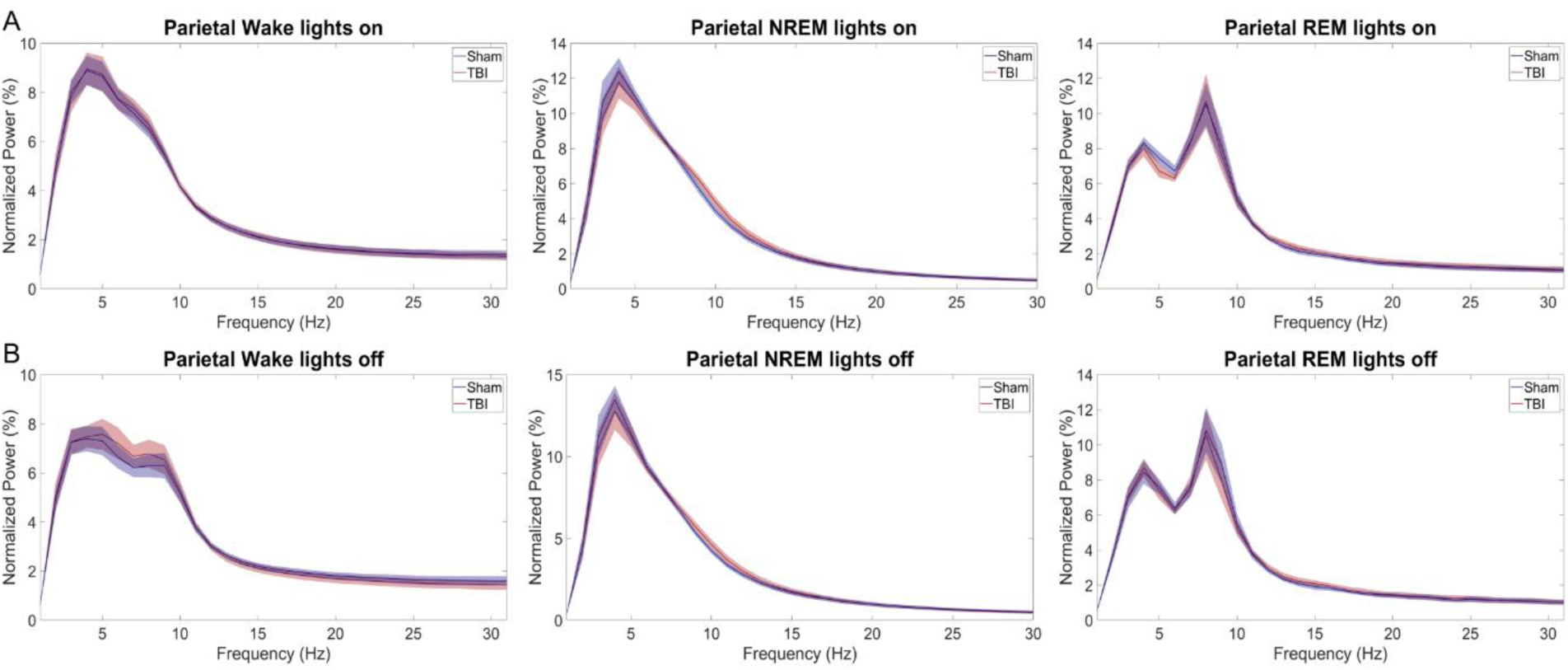
Parietal power spectral density (PSD) curves for each vigilance state during lights on and off. PSD with frequencies 0.5-30Hz in the x-axis and normalized power values in the y-axis for wake, NREM, and REM during **(A)** lights on and **(B)** lights off. Shaded area around the line represents the ±SEM. Sham data are shown in blue, and rmTBI data are shown in red. Male TBI n=6, male sham n=4; female TBI n=4, female sham n=3.

### Epileptiform activity was not increased one month post-TBI

No seizures were detected in any of the 18 animals included in this analysis. Mann-Whitney U tests revealed no significant difference in single spike or poly-spike rates (events per hour) between sham and rmTBI groups (**Fig 4A-B**). Mixed-effects model examining the interaction between treatment and vigilance state showed no significant main effects or interaction for EA rate across any vigilance state (**Fig. 4C**). A secondary exploratory analysis with sex and treatment group as independent factors similarly revealed no significant main effects or interactions for single spike or poly-spike rates (**Fig. S5A-B**). There was no significant main effect of sex or interaction effect between sex and vigilance state in the total amount of epileptiform activity **(Fig. S5C)**.

**Figure 4.**
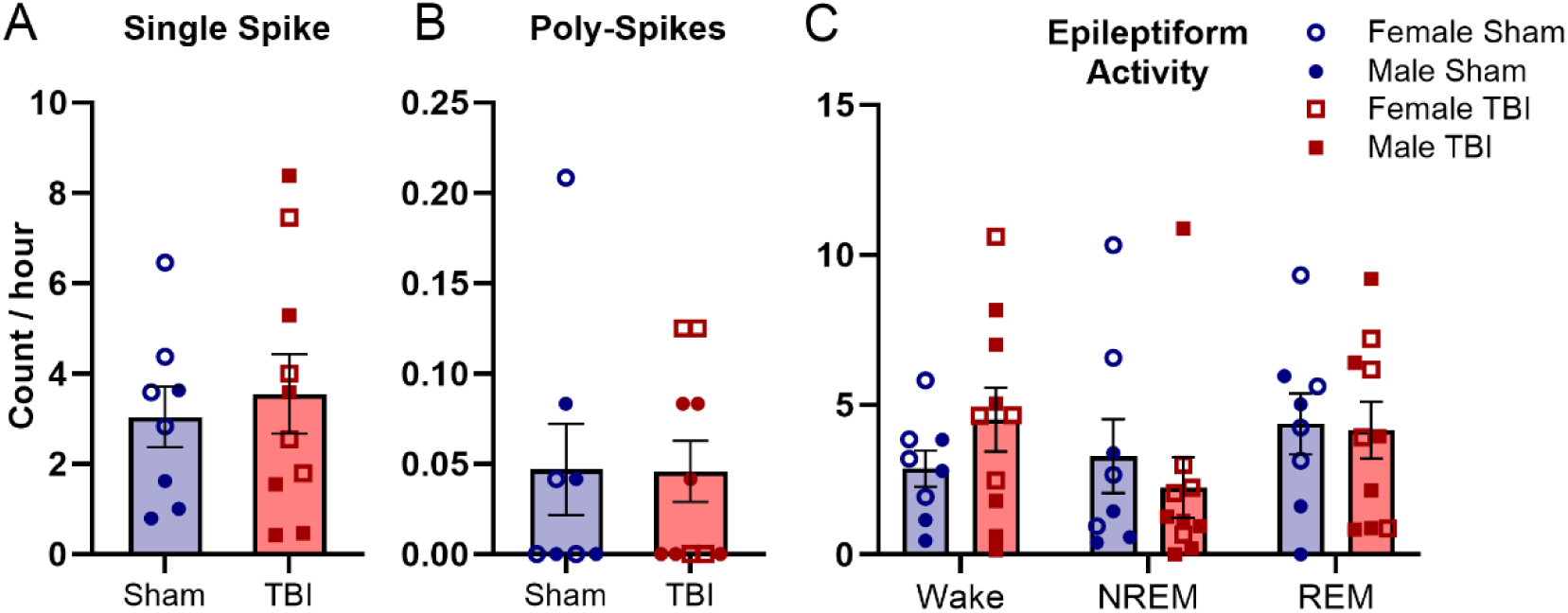
Epileptiform activity after rmTBI. **(A)** Single spike (SS) count per hour. **(B)** Poly-spikes (PS) count per hour. **(C)** Epileptiform activity (EA) count was obtained by adding SS and PS and then normalized to the total hours spent in each vigilance state. Data are reported as mean ±SEM. SS and PS counts were analyzed by Mann Whitney-U test, and EA across vigilance states was analyzed using mixed-effects model with the Geisser-Greenhouse correction. Sham data are shown in blue, and rmTBI data are shown in red. Females are shown with open symbols and males are shown with solid symbols. Each data point represents the average for a single mouse. Male TBI n=6, male sham n=4; female TBI n=4, female sham n=4.

### Neurological Injury

We assessed neurological injury by analyzing plasma samples using MSD assays to measure the levels of NF-L and GFAP with established methods (Yue et al., 2026). The levels of both plasma NF-L and GFAP from mice that received rmTBI were significantly higher relative to sham (2 folds increase in NF-L, p<0.05; 4 folds increase in GFAP p<0.05; **Fig. 5**). Two mice were excluded from analyses due to excessive contamination by hemolysis, and 1 mouse was excluded due to additional damage to the brain during EEG electrode implantation. Our findings suggest that rmTBI induces and sustains neurological injury and neuroinflammation one-month post-injury in the APP KI mouse model.

**Figure 5.**
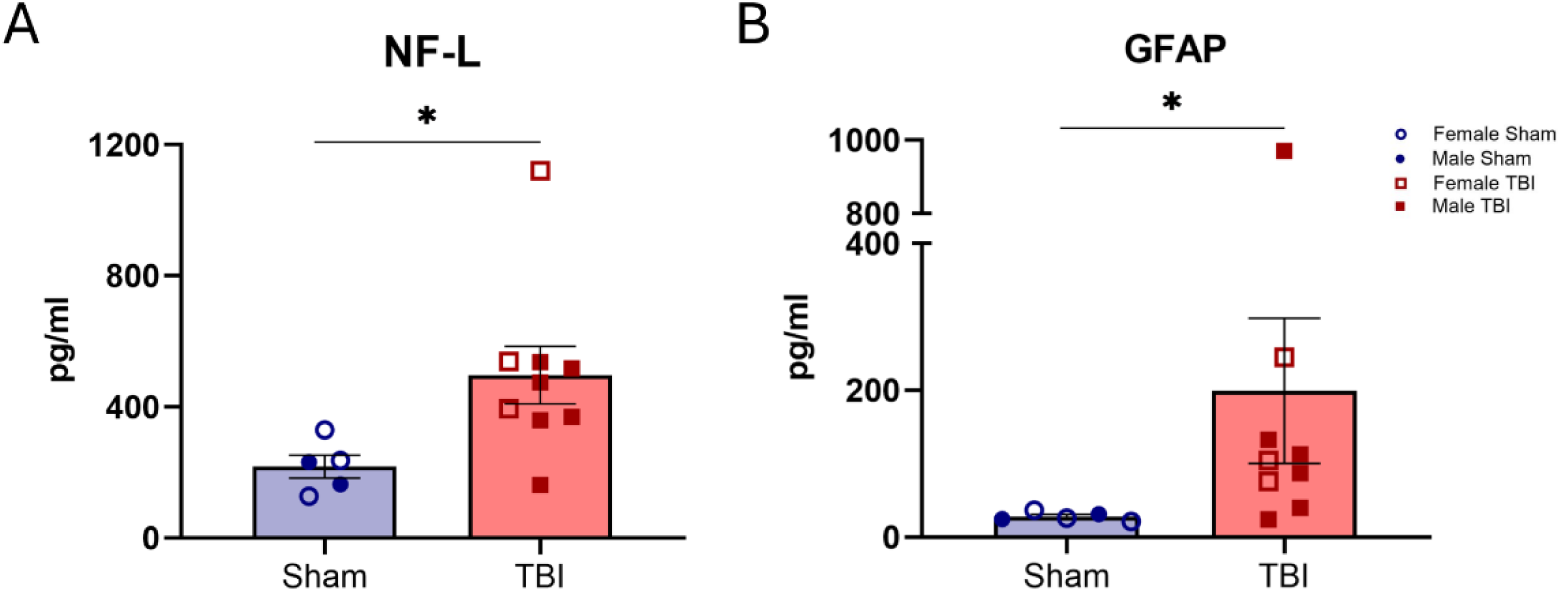
Blood biomarkers after rmTBI. **(A)** Plasma NF-L and **(B)** GFAP were compared between sham and rmTBI groups. Biomarker results were reported as mean ± SEM and analyzed by Mann Whitney-U test. Sham data are shown in blue, and rmTBI data are shown in red. Females are shown with open symbols and males are shown with solid symbols. Each data point represents a sample from a single mouse. Male TBI n=6, male sham n=2; female TBI n=3, female sham n=3. * p< 0.05.

### RmTBI associates with changes in amyloid plaque morphology

To investigate whether rmTBI affects amyloid plaque burden in the APP KI mouse model, we analyzed amyloid plaque deposition in the brain by immunohistochemistry. At the investigated age, antibody labeled plaques were only detectable in the isocortex, consistent with previous characterization of this mouse model (Masuda et al., 2016; Saito et al., 2014). There was no difference in the percentage of immunoreactive area in the isocortex between sham and rmTBI groups (**Fig 6A-B**), though a non-statistically significant uptrend in immunoreactive plaque regions could be observed in the female cohort (p=0.0754). We observed that amyloid plaques at the investigated age were clearly separated with defined circumferences. Leveraging such plaque morphologies, we extended our investigation by quantifying the number and average size of plaques in the isocortex. We found that rmTBI was associated with a downtrend in the number of plaque deposits (p=0.0513; **Fig 6D**), but plaque size demonstrated a main injury effect (p=0.0295) and an interaction effect (p=0.0010), with the female mice harboring larger plaques after rmTBI than sham (p=0.0011; **Fig 6E**). A main sex effect was also found (p=0.0489) suggesting female mice harbored larger plaques than male. These results suggest that rmTBI may alter the dynamics of amyloid plaque formation without exacerbating overall plaque burden at the onset of disease.

**Figure 6.**
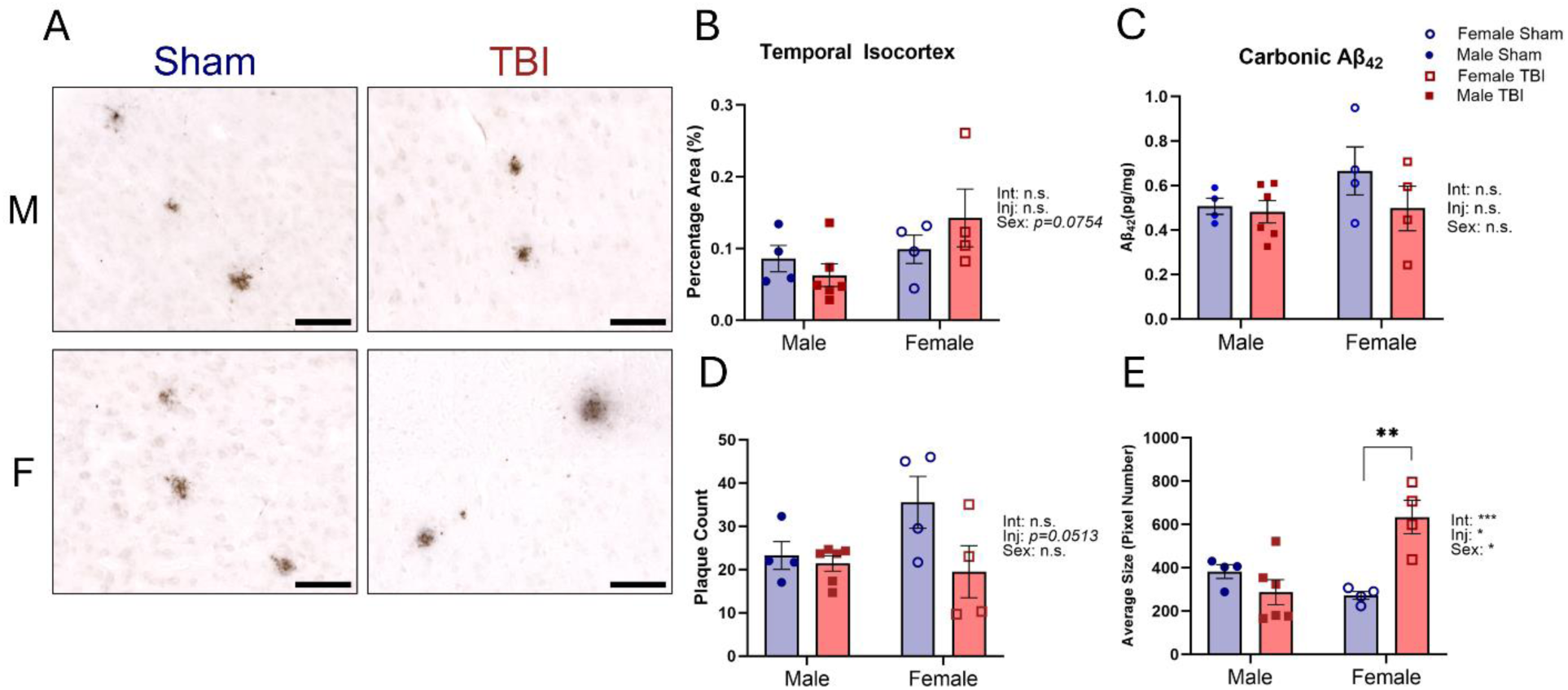
Amyloid pathology after rmTBI. Amyloid plaques were immunostained by 6E10 one month post-injury. **(A)** Representative images of the isocortex from male and female APPNL-F mice. Scale bar = 100 µm. **(B)** Quantification of the percentage area of plaques in the isocortex of both sexes. Each data point represents the average of three sections per animal. **(C)** Quantification of the carbonic (soluble) fractions of Aβ42 by ELISA using whole brain lysates. Quantification of the **(D)** average number and **(E)** average size of amyloid plaques from male and female APPNL-F mice. Each data point represents the average of three replicates per animal. Both histological and ELISA data are reported as mean ± SEM and analyzed by two-way ANOVA. Sham data are shown in blue and rmTBI data are shown in red. Females are shown with open symbols and males are shown with solid symbols. Male TBI n=6, male sham n=4; female TBI n=4, female sham n=4.

### RmTBI did not alter the level of soluble Aβ_42_ in the brain

We also assessed the levels of soluble and insoluble Aβ_42_ in hemibrain lysates by sandwich ELISA. Soluble Aβ_42_ (carbonic fraction) suggested no changes following rmTBI (p>0.05; **Fig 6C**). Insoluble Aβ_42_ (GuHCl fraction) extracted with this method did not reach detection sensitivity, likely due to the slow progression of Aβ_42_ aggregation at the investigated age. These data suggest that rmTBI did not exacerbate the accumulation of soluble Aβ_42_.

## Discussion

A major challenge in TBI research is bridging the translational gap between human and animal studies. Mild TBI in humans is highly heterogenous, resulting in diverse clinical and neurological outcomes, whereas animal studies offer controlled experimental conditions. However, rodent models of TBI vary widely and do not always recapitulate key aspects of human injury (Morganti-Kossmann et al., 2010). To address this limitation, we employed the CHIMERA model of rmTBI, which replicates human TBI biomechanics (Namjoshi et al., 2017), together with a knock-in mouse model of human AD pathology (Saito et al., 2014). We evaluated sleep architecture, PSD, epileptiform activity, and Aβ pathology one month following repeated mild TBI. We observed prolonged LRR times in the rmTBI groups, especially in females, and evidence of neuronal damage in both sexes one month following rmTBI. However, we did not find any changes in EEG sleep measures, PSD, or epileptiform activity at this timepoint post-rmTBI. Sex differences were found in vigilance state durations and PSD curves, that were independent of treatment. Female mice exhibited larger plaque sizes post-rmTBI relative to sham despite similar plaque burden.

Even though mTBI in more prevalent in males in the general population, females exhibit a greater vulnerability to persistent cognitive and somatic symptoms (Levin et al., 2021). A systematic review of mTBI reported greater symptom burden, prolonged recovery, more extensive white matter alterations, and poorer cognitive outcomes in females relative to males (Arachchi et al., 2026). Consistent with these clinical findings, we observed sex differences in loss-of-righting reflex (LRR) times following rmTBI, with female mice exhibiting longer righting latencies and increased mortality compared to males. These results may indicate greater injury severity in females at equivalent impact energy (0.7 J) or reduced recovery capacity between repeated injuries. Notably, rmTBI models are predominantly validated in male animals, with females underrepresented. Our findings highlight the need to incorporate biomarkers, such as NF-L or GFAP, to better classify injury severity and establish injury parameters that produce comparable pathophysiological outcomes across sexes.

We examined the relationship between body weight and LRR in male and female mice after observing post-TBI complications in females. An exploratory analysis revealed that females in both rmTBI and sham groups were lighter weight than males at baseline and post-injury. Female rmTBI mice lost significant body weight between baseline and the third rmTBI. In the rmTBI group, lower body weight was correlated with longer LRR times at the second and third rmTBI. Further research is needed to clarify the interactions between body weight, sex, and injury outcomes to better model injury severity across sexes.

Consistent with clinical reports of insomnia and hypersomnia persisting for days to months following mild TBI (Aoun et al., 2019; Sandsmark et al., 2017), preclinical studies have documented acute alterations in sleep architecture, including decreased NREM sleep and heightened sleep fragmentation (Morris et al., n.d.; Petraglia et al., 2014). However, the subacute effects of rmTBI on sleep in the context of AD remain understudied. In the present study, no significant differences in wake, NREM, or REM sleep duration, state transitions, or PSD were observed between sham and rmTBI groups one month post-injury, suggesting partial or full recovery of sleep/wake regulation by this timepoint. Whether this normalization reflects genuine neural recovery or transient compensation, and whether it persists over longer post-injury intervals, remains to be established. Longitudinal studies examining sleep across multiple post-injury timepoints are needed to clarify the chronic trajectory of rmTBI-induced sleep disruption and its potential contribution to long-term neurological outcomes.

Secondary exploratory analyses revealed that female mice spent more time in wake and less time in NREM relative to males, and exhibited a frontal PSD shift during wake characterized by greater low-frequency power and reduced theta-band power. These effects were independent of treatment group, suggesting they reflect underlying sex differences rather than an injury effect. This pattern is consistent with prior findings in the 5xFAD and AppNL-G-F mouse models, wherein NREM reductions relative to wild-type controls were more pronounced in females, who also exhibited greater overall sleep loss and sleep fragmentation (Kim et al., 2025; Tisdale et al., 2025). Generalized EEG slowing, characterized by increased delta power and reduced theta/alpha power, is a well-established neurophysiological hallmark of AD (Smailovic & Jelic, 2019). Collectively, the sex differences in sleep architecture observed in the present study, mainly increased wakefulness, reduced NREM, and EEG slowing in females, likely reflect a convergence of biological sex differences and AD-related pathology. The relative contributions of amyloid burden, gonadal hormonal influences, and their interaction to these sex-dependent sleep patterns warrant further investigation.

Subclinical EA occurs in early stages of AD and can contribute to cognitive decline (Vossel et al., 2017). Individuals with AD who have a seizure disorder exhibit more variable symptoms, accelerated disease progression, and brain atrophy (Lemus & Sarkis, 2023; Vossel et al., 2016). Similarly, rodent studies report increased EA in AD transgenic mice relative to wild-type controls (Diachenko et al., 2025; Gureviciene et al., 2019), and experimentally induced seizures are correlated with increased APP/Aβ expression, suggesting that EA may contribute to Aβ pathology (Kazim et al., 2017). TBI can trigger excessive glutamate release and excitotoxicity (Laskowitz & Krishnamurthy, 2016; Ng & Lee, 2019), potentially lowering the threshold for epileptiform activity (MacMullin et al., 2020) and promoting AD-related pathology.

Our data show that EA prevalence was comparable between sham and rmTBI groups, and no seizures were detected one month post-TBI. EA observed in both groups may therefore reflect the AD knock-in model rather than rmTBI effects. Consistent with this interpretation, rodent studies of PTE report seizure onset months after injury (Guo et al., 2013; Saletti et al., 2021), with incidence varying by injury severity, strain, and model (Saletti et al., 2021). Post-traumatic EA in the APP^NL-F^ mouse strain might emerge at a time point other than one month post-TBI. Additionally, our electrodes recorded at the cortical level, but since epileptiform activity has been shown to implicate deeper brain structures in AD, such as the hippocampus and entorhinal cortex, we might have missed other types of EA occurring at deeper levels, which are also early markers of hyperexcitability and can interfere with the memory system (Asadollahi et al., 2018; Hegnet et al., 2026; Kam et al., 2016; Lam et al., 2017; Soula et al., 2023). Future studies should examine long-term rmTBI effects in APP^NL-F^ mice to determine the optimal window for assessing post-traumatic epileptiform activity, and consider including microelectrodes that can travel deeper into the brain to structures like the hippocampus.

Consistent with our previous report of elevated plasma NF-L one month following rmTBI in the APP/PS1 mouse model (Yue et al., 2026), sustained neurological injury was similarly detected in APP^NL-F^ mice at this timepoint. The present injury paradigm also produced a significant increase in plasma GFAP, indicative of neuroinflammation, which was not clearly observed in the APP/PS1 model (Yue et al., 2026), likely due to the elevated baseline neuroinflammation inherent to APP-overexpressing transgenic mice. Interestingly, the female animal with the longest LRR recovery showed the highest plasma NF-L levels among female rmTBI animals. Together with our findings supporting a negative correlation between body weight and latency in LRR recovery, these findings suggest that delivering impacts at equivalent energy may produce injuries of differing severity across sexes. This highlights the importance of incorporating plasma biomarkers such as NF-L and GFAP alongside acute behavioural measures to calibrate injury intensity and ensure comparable brain injury severity between male and female animals.

The absence of significant changes in overall cortical amyloid plaque burden and soluble Aβ_42_ is consistent with our previous findings in the APP/PS1 mouse model (Yue et al., 2026). However, the observed shift toward fewer but larger plaque deposits suggests that rmTBI may alter amyloid plaque dynamics, involving various cellular and biochemical cascades not yet understood, without affecting total amyloid load. Microglia involvement in the modulation and assembly of amyloid plaques has been previously reported (Casali et al., 2020; Huang et al., 2021), and our findings raise the possibility that rmTBI stimulated glial-mediated processing of fibrillary amyloid, resulting in denser aggregates in the context of early-stage amyloid pathology. The concurrent elevations in plasma NF-L and GFAP suggest that neuronal injury and neuroinflammation occur independently of total amyloid plaque burden at this disease stage. Whether this shift in plaque morphology reflects a compensatory glial response to rmTBI or an injury-driven dysregulation of amyloid processing remains unclear. Further investigation into the responses of plaque-associated microglia and astroglia following rmTBI is warranted to elucidate the cellular mechanisms underlying these observations.

The present study has some limitations that should be considered when interpreting these findings. First, the relatively small sample size, limits the statistical power available to fully characterize sex-specific effects of rmTBI and perhaps to detect smaller effect sizes caused by the rmTBI. Second, outcomes were assessed only at a single subacute timepoint (one month post-injury), impeding any conclusions regarding the time course of rmTBI effects on sleep, epileptiform activity, or amyloid pathology. Third, since the aim of this study was to examine subacute effects of rmTBI, EEG electrodes were implanted two weeks following CHIMERA procedures, precluding the assessment of acute post-injury changes. Future studies should evaluate the immediate effects of rmTBI in the APP^NL-F^ model to determine whether this injury paradigm produces detectable changes at earlier post-injury timepoints.

Fourth, the absence of a wild-type control group raises the possibility that the lack of treatment group effects reflects a ceiling effect inherent to the APP^NL-F^ strain rather than a true absence of rmTBI-induced changes. However, exponential amyloid pathology in this strain does not emerge until approximately 12 months of age, and our histological assessment did not reveal substantial plaque accumulation at the age examined. It is therefore unlikely that strain-related ceiling effects masked rmTBI-induced changes in any of the outcomes assessed. Finally, this study did not include behavioural or cognitive assessment, which would provide a more comprehensive characterization of the functional consequences of rmTBI in this model. In addition, animals were singly housed following surgery, which may have introduced social isolation as a confounding variable, particularly for females, although this should have affected the sham and rmTBI equally, since all animals had the same housing conditions. Lastly, it is possible that three impacts were insufficient to produce detectable changes in sleep, EA, or amyloid pathology at the subacute timepoint examined. Future studies should investigate whether a dose-dependent relationship exists between the number of CHIMERA mild impacts and the magnitude of these outcomes.

## Conclusion

This study used the CHIMERA model of rmTBI in the APP^NL-F^ knock-in mouse model to examine subacute effects on sleep, epileptiform activity, and amyloid pathology. One month post-injury, rmTBI did not alter sleep architecture, PSD, or EA, despite sustained elevation in plasma NF-L and GFAP indicating ongoing neuronal injury. RmTBI did not increase total amyloid burden but shifted plaque morphology toward fewer, larger deposits. Notably, female mice showed substantially greater morbidity and mortality at equivalent impact energy, highlighting the need for sex-specific injury calibration in preclinical TBI models. These findings underscore the importance of considering sex as a biological variable in TBI research and support biomarker-guided injury standardization and longitudinal study designs in future work.

## Acknowledgement

We thank Samantha Saw, Japneet Kaur, Bonnie K. Ng, Mayuko Arai and Robert Gibson for assisting in in vivo procedures and immunohistochemistry. We thank Dr. Sarah Faber for her insightful advice on power spectra analysis. We thank the SFU animal research facility for maintaining the animals used in this study.

## Data availability

Data will be made available upon request to the corresponding author.

