## Supplemental Figures for "Repeated mild traumatic brain injury does not affect sleep or epileptiform activity one-month post-injury in a knock-in mouse model of Alzheimer’s disease"

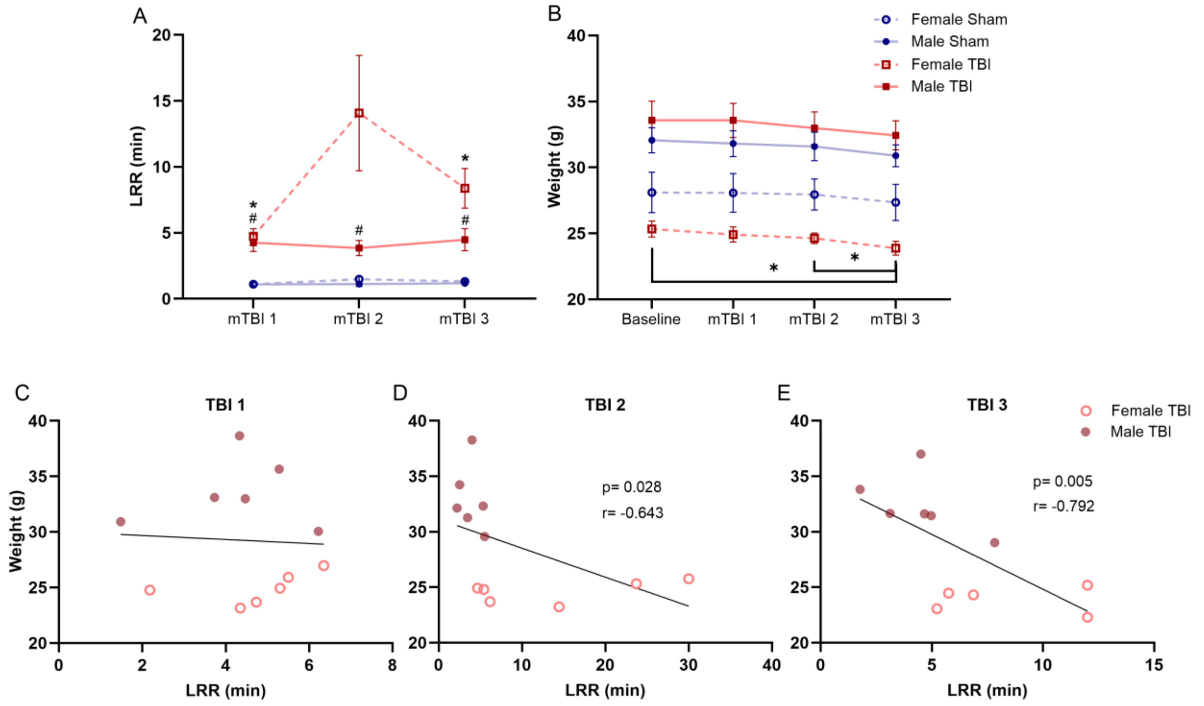


**Figure S1. Neurological behaviour and body weight after rmTBI. (A)** Average loss of righting reflex (LRR) duration in minutes for each cohort after each injury timepoint for males and females. **(B)** Body weight in grams at baseline and after each injury for males and females. Results were reported as mean ± SEM and analyzed using a mixed-effects model with the Geisser-Greenhouse correction. Sham data are shown in blue, and rmTBI data are shown in red. Males are shown with a solid line and females with a dotted line. The # symbol indicates a statistically significant difference for male sham vs male rmTBI and the * symbol for female sham vs female rmTBI. Scatter plots depict the correlation between LRR and body weight after the **(C)** first, **(D)** second, and **(E)** third mTBI. Males shown in dark red and females in light red. Females are shown with open symbols and males are shown with solid symbols. Male TBI n=6, male sham n=4; female TBI n=6, females sham n=4.

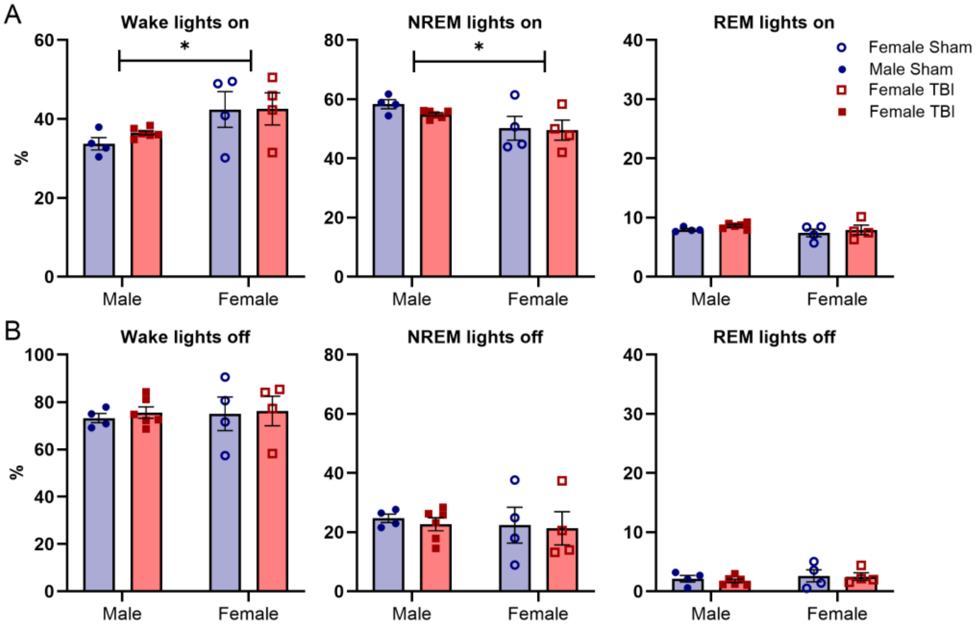


**Figure S2. Sleep architecture of APP$^{NL-F}$ mice separated by sex and light period one month after rmTBI.** Percentage of time spent in wake, NREM, and REM separated by sex during **(A)** lights on and **(B)** lights off. Data are expressed in mean ±SEM and analyzed by two-way ANOVA. Sham data are shown in blue, and rmTBI data are shown in red. Females are shown with open symbols and males are shown with solid symbols. Each dot represents the average for a single mouse. Male TBI n=6, male sham n=4; female TBI n=4, female sham n=4. * $p < 0.05$.

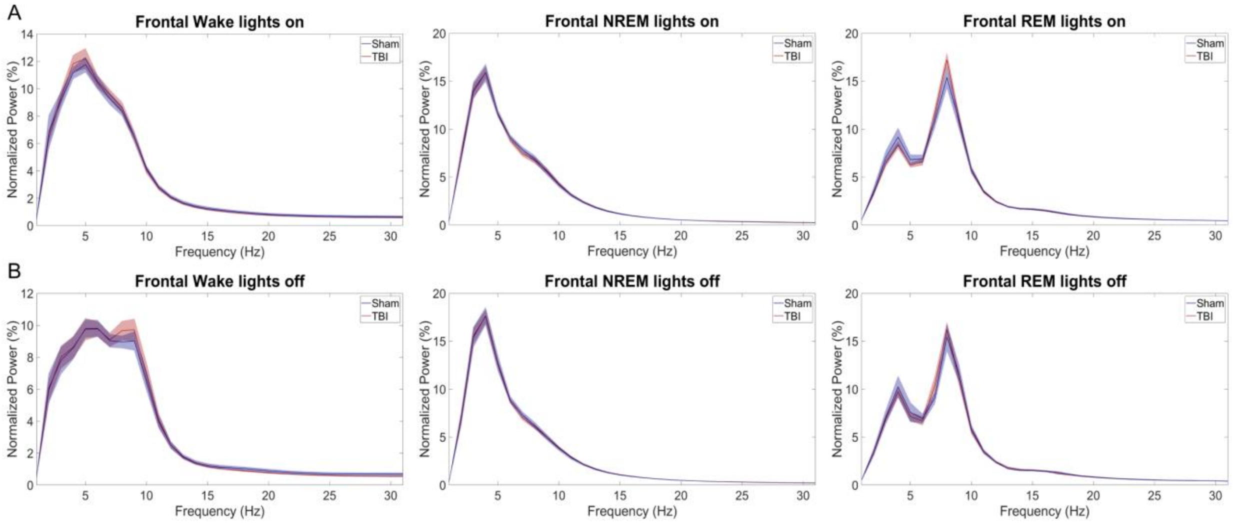

**Figure S3. Frontal power spectral density graphs across vigilance states during lights on and off**. Normalized PSD graphs for wake, NREM, and REM during **(A)** lights on and **(B)** lights off across the 24-h recording for frequencies 0.5-30Hz. Partial least squares analysis (PLS) found no differences between the group for any state or light period. Shaded areas represent the ±SEM and data were analyzed using PLS. Sham data are shown in blue, and the rmTBI data are shown in red. Male TBI n=5, male sham n=4; female TBI n=4, female sham n=4.

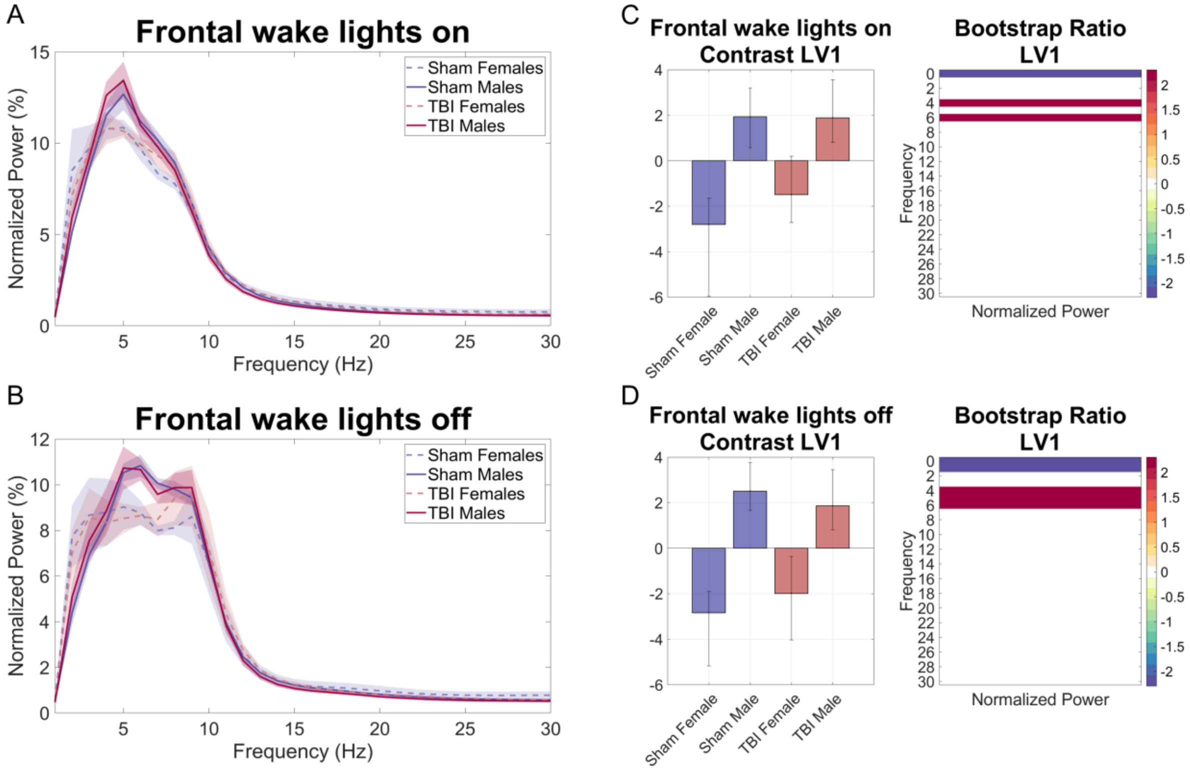


**Figure S4. Frontal wake power spectral density during lights on and off**. Normalized PSD graphs for wake during **(A)** lights on and **(B)** lights off across the 24-h recording for frequencies 0.5-30Hz. Shaded areas represent the ±SEM. Partial least squares analysis (PLS) found sex differences but no treatment group effect. **(C-D)** The PLS contrasts in male TBI, female TBI, male sham, and female sham groups during wake at the frontal electrode. Error bars represent the 95% confidence interval. The heat map represents the bootstrap ratio of the normalized power for each frequency. Purple color denotes higher power in the female groups whereas dark red color denotes lower power in the female group. Frequencies highlighted by these colors show statistical significance. Sham data are shown in blue, and the rmTBI data are shown in red. Male TBI n=5, male sham n=4; female TBI n=4, female sham n=4.

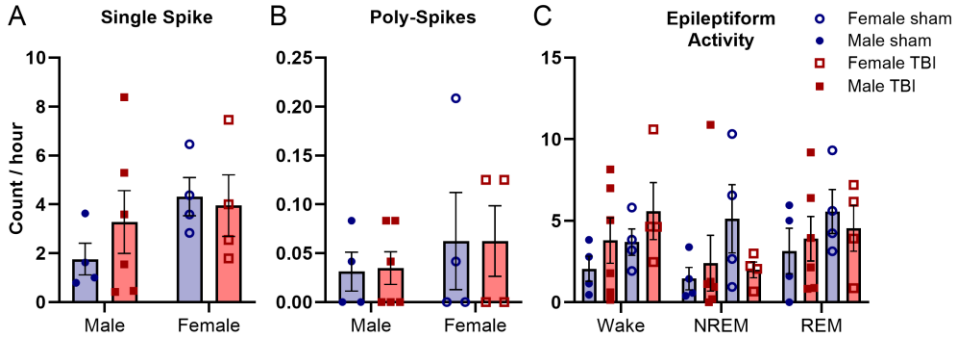


**Figure S5. Epileptiform activity one month after rmTBI.** There were no sex or TBI effects on **(A)** single spike or **(B)** poly spike counts per hour. Data are reported as mean ± SEM. Single spikes and poly-spikes were analyzed by two-way ANOVA, EA across vigilance states was analyzed by mixed-effects model with the Geisser-Greenhouse correction. Sham data are shown in blue, and rmTBI data are shown in red. Females are shown with open symbols and males are shown with solid symbols. Male TBI n=6, male sham n=4; female TBI n=4, female sham n=4.
